# Ultra-structural analysis and morphological changes during the differentiation of trophozoite to cyst in *Entamoeba invadens*

**DOI:** 10.1101/2020.06.03.131367

**Authors:** Nishant Singh, Sarah Naiyer, Sudha Bhattacharya

## Abstract

*Entamoeba Histolytica*, a pathogenic parasite, is the causative organism of amoebiasis and uses human colon to complete its life cycle. It destroys intestinal tissue leading to invasive disease. Since it does not form cyst in culture medium, a reptilian parasite *Entamoeba invadens* serves as the model system to study encystation. Detailed investigation on the mechanism of cyst formation, information on ultra-structural changes and cyst wall formation during encystation are still lacking in *E. invadens*. Here, we used electron microscopy to study the ultrastructural changes during cyst formation and showed that the increase in heterochromatin patches and deformation of nuclear shape were early events in encystation. These changes peaked at ~20h post induction, and normal nuclear morphology was restored by 72h. Two types of cellular structures were visible by 16h. One was densely stained and consisted of the cytoplasmic mass with clearly visible nucleus. The other consisted of membranous shells with large vacuoles and scant cytoplasm. The former structure developed into the mature cyst while the latter structure was lost after 20h, This study of ultra-structural changes during encystation in *E. invadens* opens up the possibilities for further investigation into the mechanisms involved in this novel process.

## Introduction

*Entamoeba histolytica*, a parasitic protist which causes amoebiasis in humans, is distributed worldwide with greater prevalence in developing countries (Debnath et al., 2012). It has simple life cycle consisting of dormant tetra-nucleate cyst which under favorable conditions converts into the actively dividing uninucleate trophozoite. These differentiation stages can be reproduced in the lab conditions in the reptilian parasite *E. invadens* since encystation has not yet been achieved in axenically grown *E. histolytica* trophozoites (Sanchez et al., 1994). In encystation process of *E. invadens* motile trophozoite rounds up and eventually becomes encapsulated within a rigid, relatively impermeable cell wall (Chavez et al., 1978). The cyst wall components are concentrated and transported in specific encystation specific vesicles (ESV) which contain a fibrillar material (Chavez-Munguia et al., 2003). This material is digested on the plasma membrane through the rupture of the vesicles. Similar type of ESVs are also found in *Acanthamoeba castellanii* (Chavez-Munguia et al., 2005) and *Giardia* (Reiner et al., 1990). This morphological evidence supports that the vesicles found in the amoeba and those identified in the flagellated *G. lamblia* may correspond to a common biosynthetic mechanism of synthesis, transport and deposition of cell wall components (Chavez-Munguia et al., 2008). Literature indicates that *Entamoeba* undergoes sexual or parasexual reproduction at some stage but timing is not known (Ali et al., 2008, Ramesh et al., 2005, Weedall et al., 2012, Weedall and Hall, 2015). Meiosis specific genes were found upregulated and homologous recombination was observed during stage conversion or under stress conditions (Singh et al., 2013).

During encystation, nuclear division takes place in the absence of cytokinesis, such that the uninucleated trophozoite transforms into a dormant tetra-nucleated cyst, enveloped by a protective cyst wall. Detailed studies on the mechanism of cyst formation, information on ultra-structural changes during encystation are lacking in *E. invadens*. In this study, we attempt to address the mechanism of cyst formation by transmission electron microscopy (TEM) and fluorescence microscopy.

## Results

### Nuclear dynamics during encystation

Nuclear shape and heterochromatin (HC) distribution of *E. invadens* was investigated by transmission electron microscopy during encystation. In normal proliferating trophozoite, condensed HC masses are arranged directly underneath the nuclear membrane and the nuclear shape is spherical (Fig. 1) as reported previously in *E. histolytica* (Chavez-Munguia et al., 2008, Dahl et al., 2008, Solis and Barrios, 1991). HC is densely packed chromatin, which usually reflects modifications of DNA, histones and other DNA binding proteins, and is typically transcriptionally inactive (John, 1988). In *Entamoeba*, peripheral HC contains the ribosomal DNA circles and stains with nucleolar markers like fibrillarin (Jhingan et al., 2009). We looked for changes in nuclear distribution of HC during encystation and found that large patches of HC began to appear as early as 4h after induction of cyst formation, and by 20h the nuclei in 90% cells had these patches of HC (Fig. 1, 3). At 24h and 48h, HC patches were seen not only at the periphery but at internal locations as well. The nuclear shape began to change from spherical to distorted within 2h of encystation, and by 16h, ~95% cells had distorted nuclei. By 20h most cells had nuclei with HC patches and distorted shape (Fig. 3), after which there was a gradual increase in nuclei with the normal pattern. In mature cysts (72h) the normal pattern of HC and nuclear shape was restored (Fig. 1). Chromatoid bodies composed of poly ribosomes appeared in the cytoplasm after 16h.

**Figure 1:**
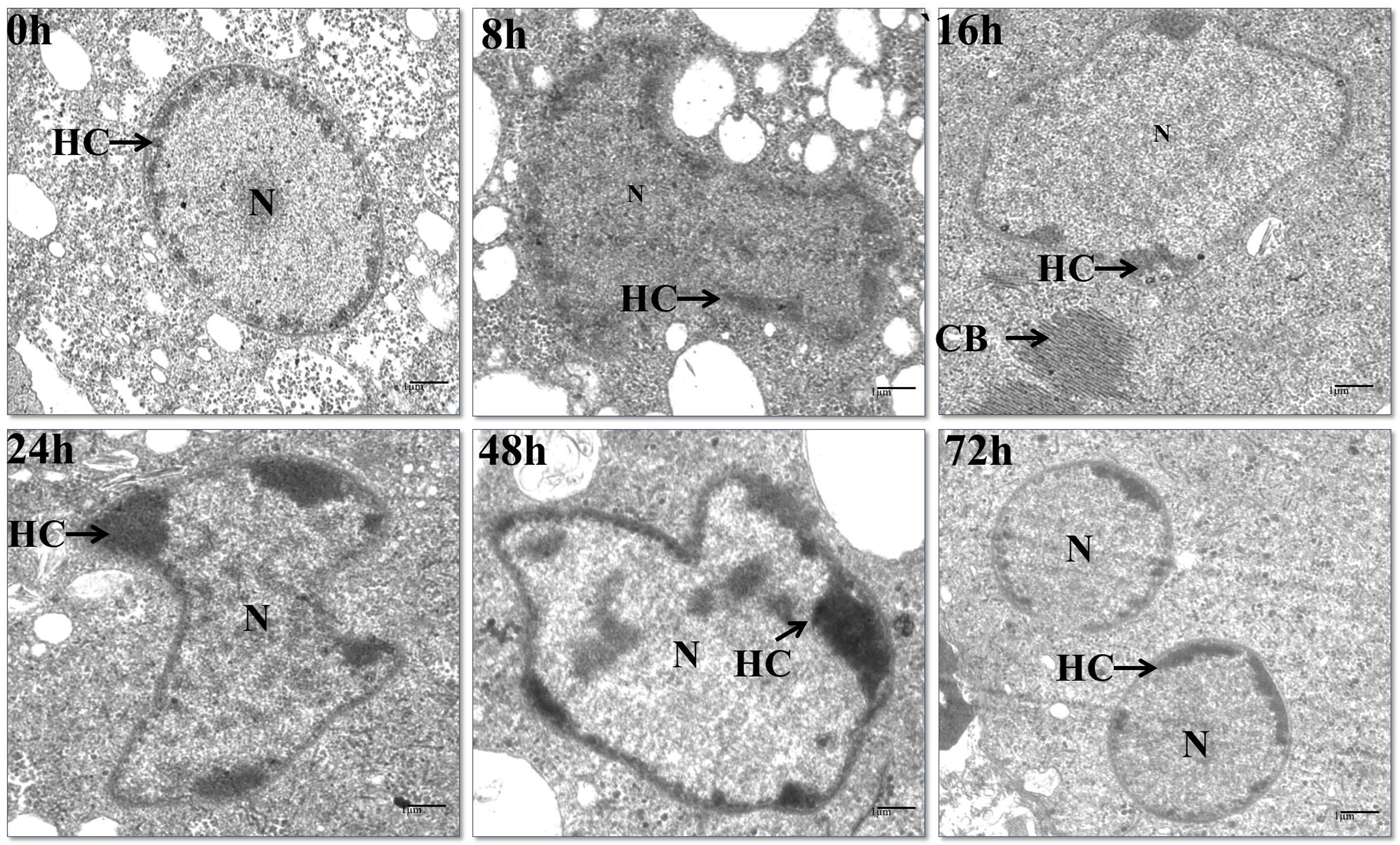
Transmission electron micrograph showing the changes in the nuclear structure and heterochromatin distribution in *E. invadens* during encystation. Samples were taken at different time points i.e. 0h, 8h, 16h, 24h, 48h, and 72h, after transfer to LG medium. HC; heterochromatin, N; nucleus, CB; chromatoid body.

**Figure 2:**
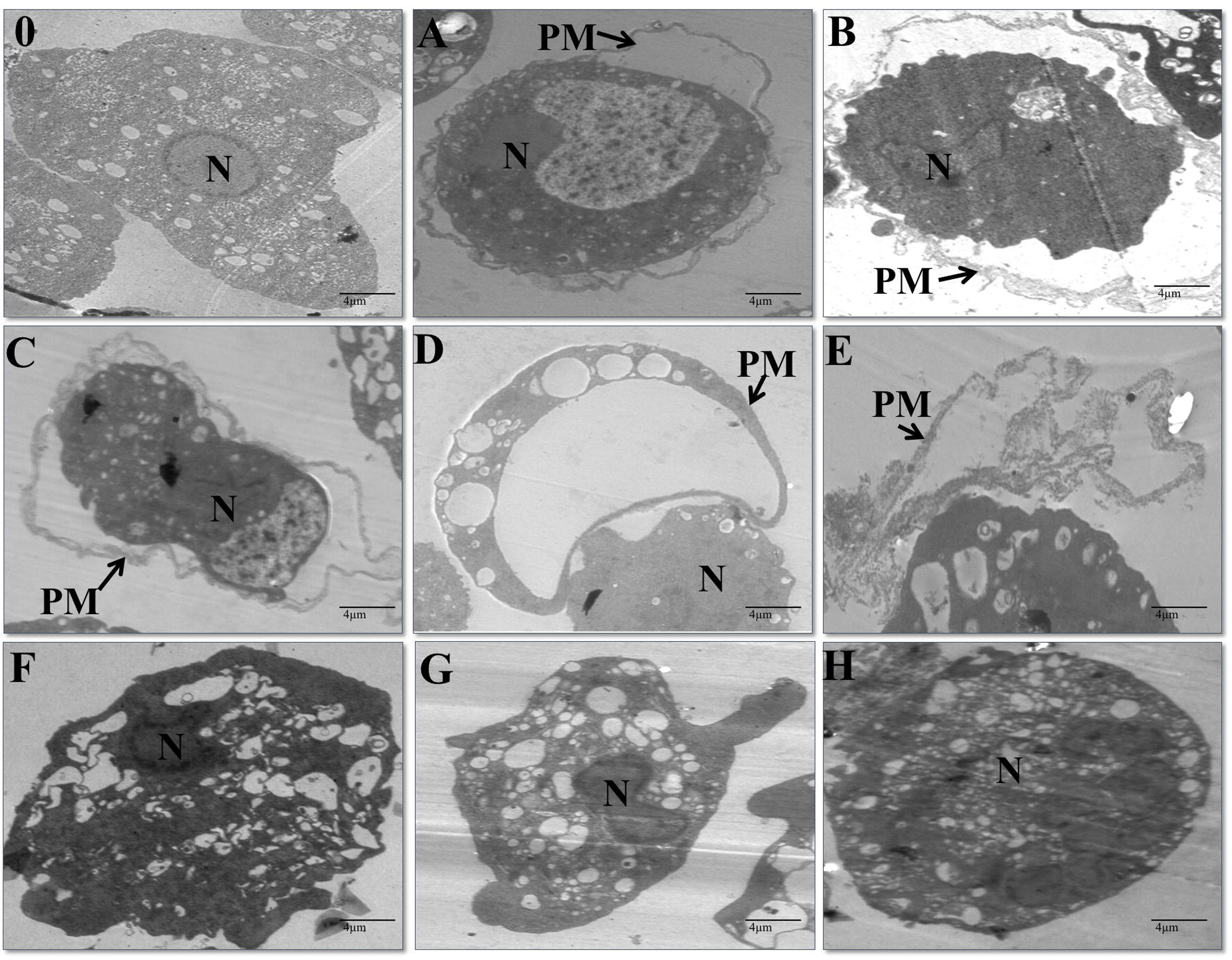
Formation of cyst within plasma membrane showed by electron microscopy in *E. invadens* during encystation. 0; normal proliferating trophozoite, A; 8h, B, C; 12h, D; 16h, E; 20h, F; 24h, G; 48h, H; 72h. N; Nucleus, PM; Plasma membrane.

**Figure 3:**
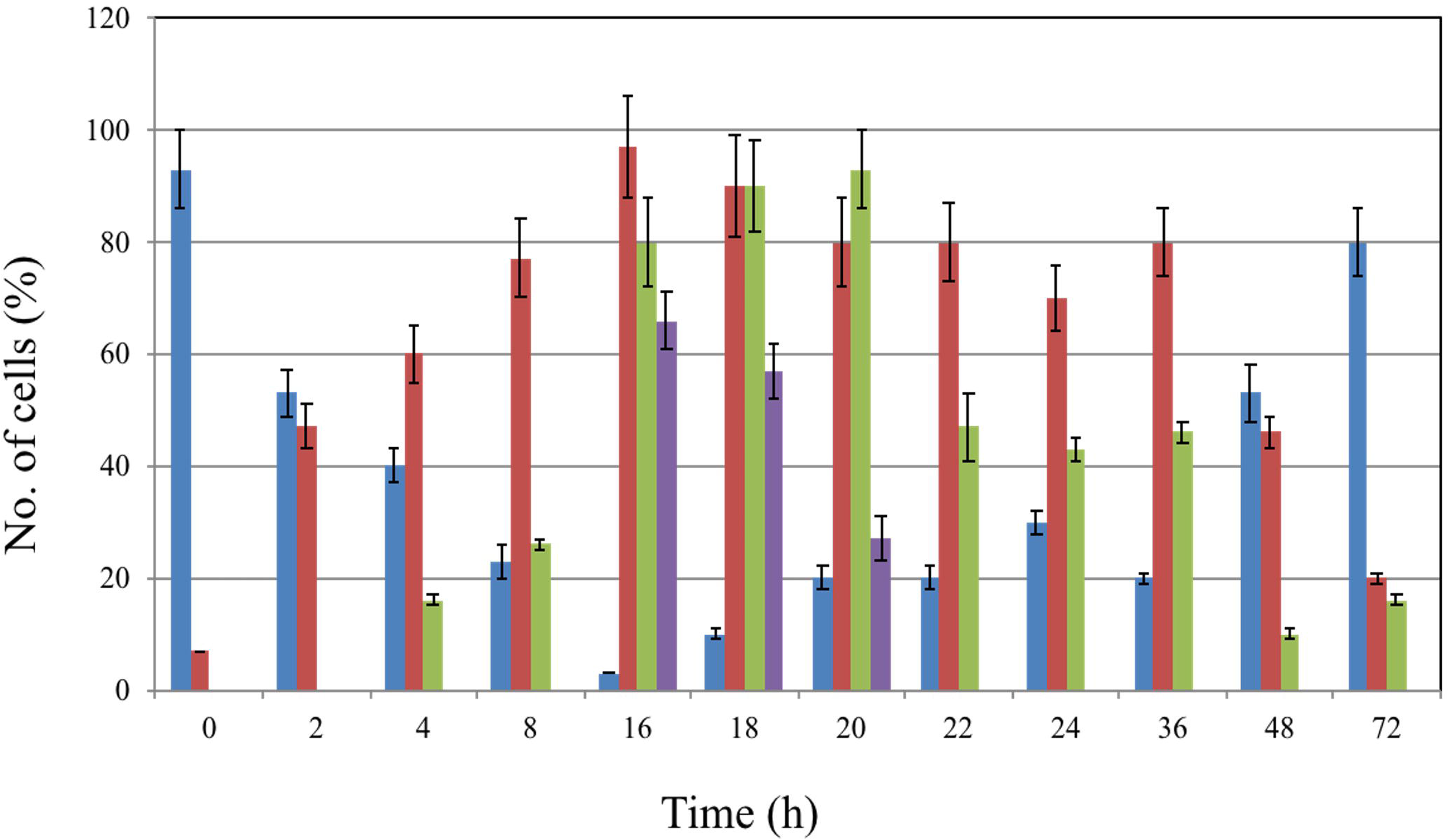
Distribution of different cell type during encystation in *E. invadens* investigated by electron microscopy. Blue; normal nucleus, Brown; distorted nucleus, green; heterochromatin patches, purple; detached plasma membrane.

### Morphological changes during cyst formation

Sequential ultra-structural changes occurring in the trophozoite during encystation were investigated through TEM. Trophozoites were inoculated in encystation medium and samples were fixed after each time point. Thin sections were examined for morphological changes between 8h to 16h (Fig. 2). In high resolution electron microscopy at zero time point, trophozoites showed normal circular nucleus and no chromatoid bodies. Plasma membrane was in uniform contact with cytoplasm (Fig. 2, 0h). Remarkable changes were visible after 8h. In a number of cells the cytoplasmic mass could be seen separated from the plasma membrane (Fig. 2A). Gradually the separation between plasma membrane and cytoplasmic material increased, and in some cells the cytoplasmic mass was almost completely detached from the membrane (Fig. 2, A-C). By 16h, ~60% cells showed a near complete separation of the plasma membrane and cytoplasm (Fig. 3). Two types of structures were visible. One was densely stained and consisted of the cytoplasmic mass with clearly visible nucleus. The other consisted of membranous shells with large vacuoles and scant cytoplasm (Fig. 2, D-E). After 20h the latter were lost and the former developed into mature cyst, with the spherical nuclei being restored at 72h.

We also observed the encysting cells in real time to further understand how the above mentioned phenomenon took place. Cells were inoculated in encystation medium and monitored under confocal microscope between 8h to 12h. It is well documented that cells which are getting ready to encyst undergo clump formation and clumping is a prerequisite for encystation. Thus it was not possible to focus on well-separated cells and follow them in real time. In addition, there was rapid pseudopodial movement and constant movement of neighboring cells in and out of the viewing field. Due to these reasons it was difficult to observe a single encysting cell in a sequential manner over an extended time period. Our observations of cells 8-12 h post-induction (Fig. 4) show that cells (marked by arrow in Fig. 4A) in a clump of trophozoites could be seen rounded and with a clear zone in the center, with some cytoplasmic material on the periphery. With time, structures became visible which had large vacuoles with little cytoplasmic material and looked like hollow barrels or cups (marked by * in Fig. 4 F, G, J, K) symbol marked (Mam).(marked by O in Fig. 4 B, C, D, E, G, H, I, J, K). Simultaneously, one could also see the accumulation of round cytoplasmic masses (with almost the same diameter as the opening of the barrel/cup), these are marked by ▲ in Fig. 4 B, C, D, F, H-N. Whether the cytoplasmic spheres are extruded from the vacuolated cell or the two are independent entities needs to be resolved. At later times (panels M-P) very few original trophozoites are visible, and the structures seen are either the transparent vacuolated shells or the dense cytoplasmic spheres.

**Figure 4:**
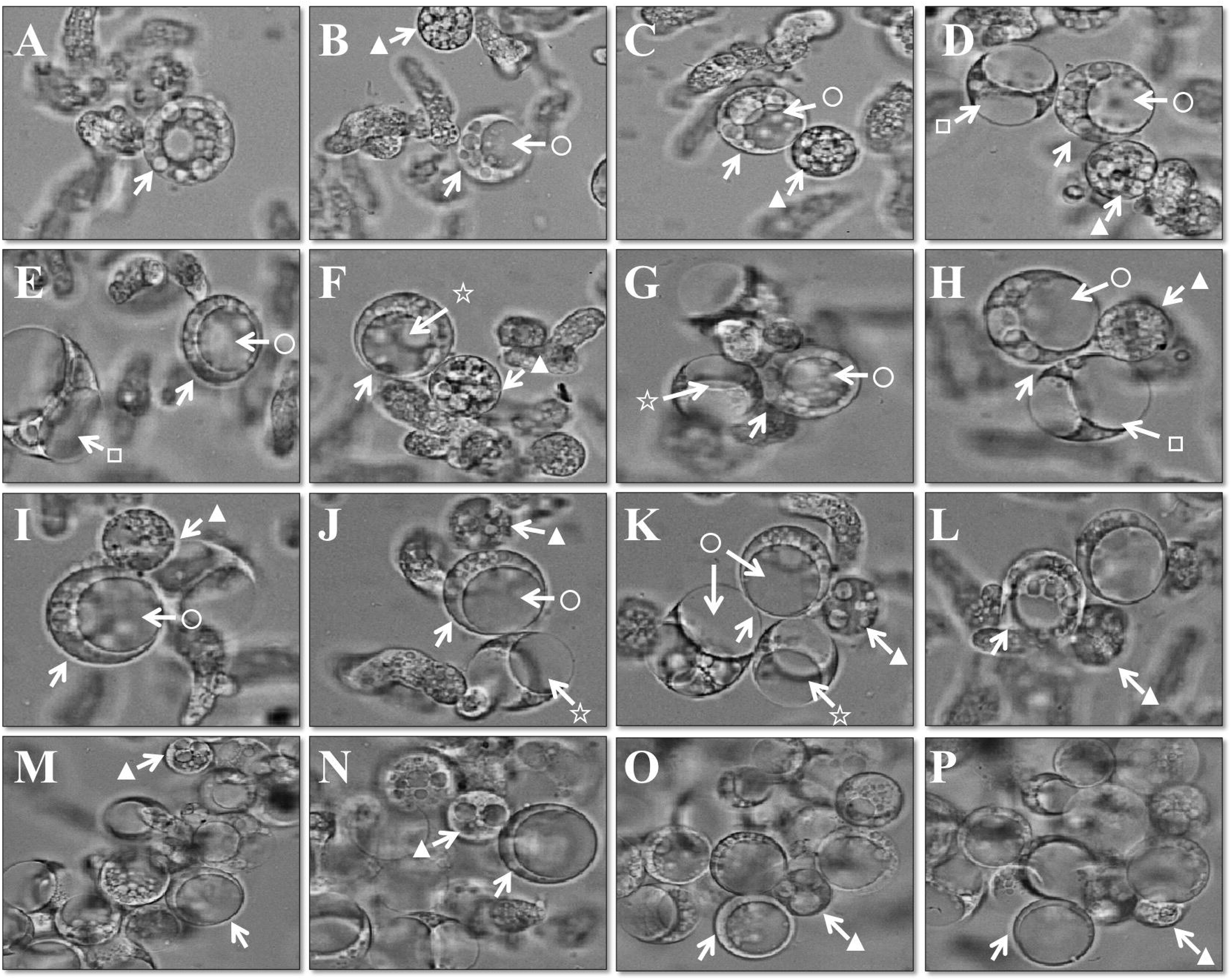
Live cell imaging of encysting *E. invadens* trophozoites. Cells were inoculated into induction medium and viewed by microscopy between 8 to 12h. Arrow indicated the plasma membrane of particular cell (ghost).

That the vacuolated shells were indeed plasma membrane bound was shown with a plasma membrane specific dye (“cell mask” red cat no. C10046 from Invitrogen) which specifically binds to plasma membrane. Membrane staining was also done with peroxiredoxin antibody which has been shown to localize preferentially to the plasma membrane in *Entamoeba* (Choi et al., 2005). In normal trophozoite the membrane specific reagents labeled the membrane, with some internal diffusion (Fig. 5). The 16h sample showed very clear labeling of the vacuolated shell with “cell mask” dye and the absence of nucleus was confirmed by lack of Hoechst staining. The dense cytoplasmic spheres were uni-nucleate. Peroxiredoxin antibody also showed the same pattern in the vacuolated shell, which lacked any nucleus.

**Figure 5:**
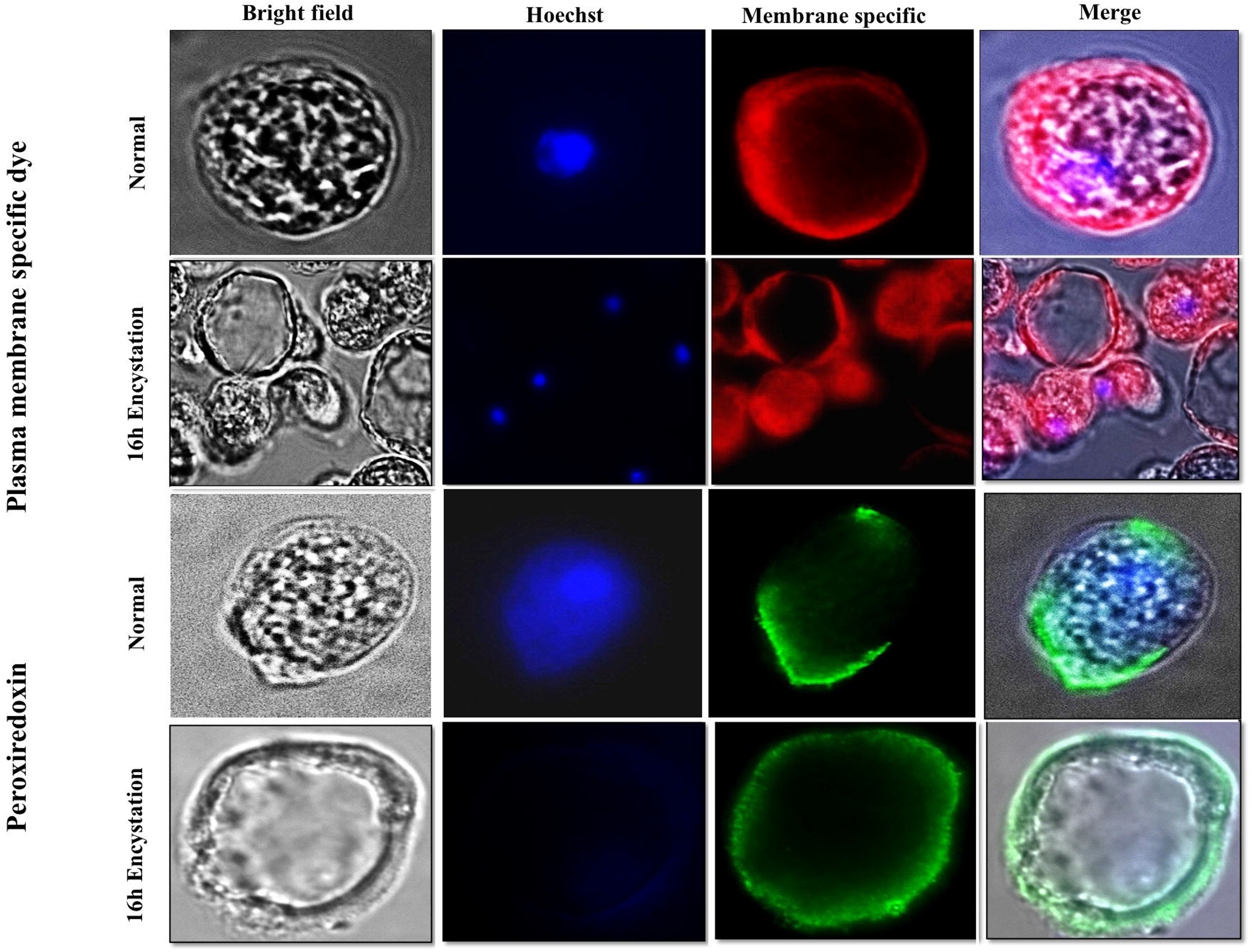
Staining with plasma membrane specific reagents. Cells were taken at 16h during encystation. Peroxiredoxin antibody was kind gift from Sharon Reed (Choi et al., 2005) which localized to plasma membrane of *E. invadens*. Green; anti mouse-peroxiredoxin primary antibody (1:250) and anti-mouse-secondary antibody tagged with alexa-488 (1:500), red; plasma membrane specific red dye “cell mask” cat no. C10046 from Invitrogen. Hoechst 33342 was used as nuclear stain.

## Discussion

Our EM study showed that the nucleus of *E. invadens* in normal proliferative cells was typically spherical or ellipsoid which was identical to *E. histolytica* (Chavez-Munguia et al., 2008, Solis and Barrios, 1991). Heterochromatin was present at the periphery, which is also the site of nucleolar localization (Jhingan et al., 2009). In this respect also *E. invadens* is identical with *E. histolytica*. HC is generally associated with low activity of gene expression, and many specific proteins and characteristic histone modifications present in heterochromatin are responsible for gene silencing (Dahl et al., 2008, Sarma and Reinberg, 2005). Conversely, euchromatin is rich in transcriptionally active genes and is often located at the nuclear interior in more open chromatin structures. In *E. invadens*, the nuclear shape changed and larger patches of HC were visible during encystation (Fig. 1). Our results showed that the increase in HC patches was an early event during encystation. It could be related to up- or down regulation of stage specific genes which takes place in cells committed for encystation (Ehrenkaufer et al., 2013, Manna et al., 2014).

It has been observed that open euchromatin structures are more deformable than tightly packed heterochromatin structures (Pajerowski et al., 2007), and any external or intracellular forces could reorganize gene rich areas relatively easily. Even though the nucleus is the stiffest cellular organelle and is 2- to 10-times stiffer than the surrounding cytoskeleton, (Caille et al., 2002, Guilak et al., 2000), extracellular forces and strain result in clearly detectable nuclear deformations (Guilak, 1995, Guilak et al., 2000, Maniotis et al., 1997). An abnormal nuclear shape is frequently associated with cancer (Webster et al., 2009, Zink et al., 2004). It has been speculated that change in nuclear shape leads to change in chromosome organization which in turn can alter gene expression (He et al., 2008). Altered nuclear shape in cancer cells is thought to facilitate metastasis because of reduced nuclear stiffness, which could increase the ability of transformed cells to penetrate tissue (Dahl et al., 2008). Inactivation of some proteins which associate with endoplasmic reticulum can affect nuclear shape (Higashio et al., 2000, Matynia et al., 2002). Nuclear shape can also be affected by lipid synthesis. This has been shown in yeast and *Caenorhabditis elegans*, where the inactivation of a lipid phosphatase that is homologous to the mammalian lipin (Reue and Zhang, 2008) was shown to cause expansion of the ER membrane and alteration in nuclear shape (Campbell et al., 2006, Golden et al., 2009).

Changes in nuclear structure and function could contribute both to increased cellular sensitivity to stress and to altered transcriptional regulation (Lammerding et al., 2004). Some specialized cells undergo dramatic changes in nuclear shape during differentiation and maturation. For example, spermatids have extremely elongated nuclei (Dadoune, 2003) and neutrophils develop extremely lobulated nuclei, which are associated with loss of lamin A/C (Yabuki et al., 1999) and expression of lamin B receptor (Hoffmann et al., 2007). Thus changes in nuclear shape are well-documented with respect to cellular differentiation, stress and disease. We speculate that adaptations in nuclear shape and structure are directly related to the functionality of the *E. invadens* cell and in the differentiation process.

In this study, we also documented the detailed observation of ultra-structural changes during encystation in *E. invadens*. Encystation is similar to the well-known process of sporulation in yeast and bacteria in that both are triggered by adverse growth conditions, and result in production of a resting cell that is more resistant to environmental factors than the vegetative cell. In *S. cerevisiae* during sporulation, a meiotic division takes place, resulting in the production of four haploid nuclei which are then enveloped by de novo formed plasma membranes within the cytoplasm of the mother cell to form immature spores or prospores (Neiman, 2011). In *Bacillus subtilis*, a single spore (fore spore) is formed by a process of engulfment following an asymmetrical division within the mother cell. Upon completion of spore formation the mother cell lyses, releasing the mature spore (Higgins and Dworkin, 2012).

The major difference between the trophozoite and cyst in *E. invadens* is that the latter is about half the average diameter of the former, and while the trophozoite is uni-nucleate, the cyst is tetra nucleate. In addition, the cyst possesses a chitin wall. Since one trophozoite gives rise to only one cyst it was assumed that once the differentiation process is set in motion in *E. invadens*, the process would be relatively straightforward. Our TEM and confocal observations reveal a more complex picture. We find that once the initial signal for differentiation is conveyed to the trophozoite, the cytoplasmic mass begins to move away from the original plasma membrane. At this stage the spherical cytoplasmic mass may be referred to as a pre-cyst and the chitin wall is not yet visible by calcofluor staining. In addition, it contains only one nucleus. A second class of structures is visible which consists of large vacuoles bound by plasma membrane and containing little cytoplasm. Whether the pre-cyst is extruded as a spherical mass, leaving behind this empty shell, or the two structures are unrelated needs to be understood. It is possible that the vacuolated structures represent cells that have undergone apoptosis and would not contribute to cyst formation. An altruistic possibility may also exist whereby the death of these cells could benefit other cells to complete the differentiation process (Picazarri et al., 2008). In a study of autophagosome-like structures in encysting *E. invadens* cells it was shown that the autophagy gene, Atg8 was associated with structures that looked like vesicles/vacuoles as early as 2 h after induction of encystation. Some of these had diameter >4 μm, and they peaked at 24 h when >65% of cells contained these structures (Picazarri et al., 2008). It is possible that these could be the precursors of large vacuolated cells reported in our study.

Our data show that *E. invadens* has evolved a unique process of cyst formation, the details of which are not clearly understood. This work opens up the possibilities for further investigation into the mechanisms involved, which might help to understand the significance of this novel process.

## Materials and Methods Strains and cell culture

All experiments were carried out with *E.invadens* strain IP-1 was obtained from the American Type Culture Collection and maintained at 25°C in TYI-S-33 containing 15% heat inactivated adult bovine serum,125ml/100 mlstreptomycin/penicillin G and 2.0% vitamin mix (Singh et al., 2012).

### Cyst induction and excystation

This was done essentially as described and adapted in our lab (Singh et al., 2011). Briefly, log phase trophozoites grown in 50 ml flasks were chilled on ice for 10 min to remove the cells from the wall and harvested by centrifugation at 500g for 5 min at 4°C. 5×10^5^ trophozoites ml^-1^ were transferred into induction medium (LG): TYI medium was prepared without glucose and diluted to 2.12 times with water and completed with 5% heat inactivated adult bovine serum, 2.6% vitamin mix and 125 mL/100 ml antibiotic. Cysts obtained in LG medium after three days were harvested and treated with 0.05% Sarkosyl (Sigma) to destroy the trophozoites. Typical spherical refractile cysts were observed by light microscopy and checked for staining with calcofluor and observed 80–90% encystation efficiency. For excystation, the cysts were further washed with phosphate-buffered saline (PBS), and inoculated in normal TYI-S-33 medium.

### Electron microscopy

To examine cellular and nuclear morphology during encystation, *E. invadens* cells were grown till late log phase and 5 x 10^5^ cell/ml was transfer into LG medium for cyst induction for different time points and fixed in 2.5% glutaraldehyde2.0% paraformaldehyde in 0.1M PBS, pH 7.2 for 2h at RT and overnight at 4°C. Washed thrice with 0.1M PBS. Specimens were post fixed in 1% osmium tetroxide and 0.1M PBS, for 2h at 4°C, washed with 0.1M PBS. The specimens were dehydrated with increasing concentrations of ethanol and then clearing for 2h in toluene at RT, followed by infiltrate in a 1:1 mixture of toluene and Epoxy resin-Araldite (M) EY212 (TAAB laboratory equipment ltd.) overnight. Specimens were subsequently embedded in Epoxy resin at 60°C for 2 days. Semi- and ultra-thin sections were obtained using an ultra-microtome Leica UC-6. Semi-thin sections (1μ) were dried on slides at 80°C and stained with a mixture of 1.0% methylene blue. Ultra-thin section (70-90nm) on 200 mess Copper grids stained with saturated solution of uranyl acetate in 50% alcohol for 15 min followed by lead citrate (50 mg/12ml MQ) for 10 min at RT. Sections were scanned and photographs were taken with a JEOL-JEM-2100F transmission electron microscope at 120 kV and scanned images were processed using Adobe Photoshop software. Sample preparation and viewing was done at Advanced Instrumentation Research Facility (AIRF), Jawaharlal Nehru University, New Delhi, India.

### Immuno-fluorescence microscopy

Cells were washed with PBS, fixed in cold methanol for 20 min at −20°C, again washed twice with PBS buffer and permeabilized with 0.1% triton-X-100 for 5 min at room temperature, washed and blocked with 3% BSA at RT for 60 min. After blocking cells were washed once with PBS and incubated for 45 min at RT in monoclonal/polyclonal antibody generated in mouse/rabbit in suitable dilutions, washed with PBS and incubated for 30 min with anti-mouse/rabbit secondary antibody tagged with fluorochrome in suitable dilutions along with nuclear strain Hoechst 33342 (2.5 μg/ml). For positive control, polyclonal anti-rabbit/mouse anti-fibrillarin antibody (Jhingan et al., 2009) was used in each preparation in 1:500 dilutions. Trophozoites were then washed thrice with PBS and mounted on glass slide in 25% glycerol with anti-fed p-phenylenediamine (sigma). All steps were done in 1.5ml microfuge tubes, and centrifuged at 4°C; 500g for 5 min. Phase contrast and fluorescent images were taken using a Zeiss Axio Imager.M1 microscope (Germany) with a 100x magnification

## Acknowledgement

NS and SN were recipient of fellowship from CSIR, India. SB is a recipient of J.C. Bose fellowship. We thank Prof. Alok Bhattacharya, School of Life Sciences Jawaharlal Nehru University for his critical inputs and scientific discussions. We also thank Advanced Instrumentation Research Facility (AIRF) and Dr. Gajender Saini, Jawaharlal Nehru University for his help during sample preparation and Electron Microscopy.

## Notes

### Competing Interest Statement

The authors have declared no competing interest.

